## Supplementary material for "WRNIP1 prevents transcription-associated genomic instability": Suppl information

### APPENDIX MATERIALS AND METHODS

#### Immunofluorescence

Immunofluorescence for γ-H2AX was performed as previously described (Murfuni et al. 2012). Briefly, exponential growing cells were seeded onto Petri dishes, then treated (or mock-treated) as indicated. Subsequently, cells were fixed in 4% formaldehyde for 10 min, permeabilized using 0.4% Triton X-100 for 10 min and incubated with 10% FBS for 1 h at RT. After blocking, for γ-H2AX detection, cells were incubated with the mouse-monoclonal anti-γ-H2AX primary antibody (Millipore; 1:1000) for 1h at RT.

FANCD2 immunostaining was performed similarly to γ-H2AX, with the exception that cells were pre-extracted with ice-cold CSK buffer for 10 min on ice. After blocking, cells were incubated with the primary anti-FANCD2 antibody (Santa Cruz Biotecnology; 1:250) for 1 h at RT.

Following each primary antibody incubation, cells were washed twice with PBS and then incubated with the specific secondary antibody: goat anti-mouse Alexa Fluor-488 or goat anti-rabbit Alexa Fluor-594 (Molecular Probes). Secondary antibody incubation was carried out in a humidified chamber for 1 h at RT. DNA was counterstained with 0.5 μg/ml DAPI.

Images were randomly acquired using an Eclipse 80i Nikon Fluorescence Microscope equipped with a Video Confocal (ViCo) system. For each time point, a minimum of 200 nuclei were examined. Nuclear foci were scored at a 40× magnification, and only nuclei showing more than five bright foci were considered positive. Intensity per nucleus was calculated using ImageJ. Parallel samples incubated with either the appropriate normal serum or only the secondary antibody confirmed that the observed fluorescence pattern was not attributable to artefacts.

#### Chemicals

The transcription elongation inhibitor Cordycepin (Cordy; Sigma-Aldrich) was employed at a final concentration of 50 µM. The stock solution was prepared in DMSO at a concentration >1000. The final concentration of DMSO in the culture medium was consistently kept below 0.1%.

#### RNA interference

The RAD18 genetic knockdown experiment was performed using Interferin (Polyplus) following the manufacturer's instructions. A Flexitube siRNA (QIAGEN) was utilized, targeting the following sequence of the mRNA (5'-ATGGTTGTTGCCCGAGGTTAA-3'), with the siRNA used at a concentration of 10 nM. As a control, a siRNA duplex directed against GFP was employed. Depletion was confirmed by Western blot using the relevant antibodies (*see below*).

**Western blot analysis**

The proteins were separated on polyacrylamide gels and transferred onto a nitrocellulose membrane using the Trans-Blot Turbo Transfer System (Bio-Rad). Subsequently, the membranes were blocked with 5% NFDM in TBST (50 mM Tris/HCl pH 8.0, 150 mM NaCl, 0.1% Tween-20) and incubated with the primary antibody for 1 h at RT. The primary antibodies employed for Western blot were rabbit-polyclonal anti-RAD18 (Abcam; 1:2000) and rabbit-polyclonal anti-LAMIN B1 (Abcam; 1:30000). Following the primary antibody incubation, the membranes were probed with horseradish peroxidase-conjugated goat specie-specific secondary antibodies (Santa Cruz Biotechnology; 1:20000) for 1 h at RT. Signal visualisation was achieved using Western Bright ECL HRP substrate (Advansta) and captured with Chemidoc XSR+ (Chemidoc Imaging Systems, Bio-Rad).

#### APPENDIX FIGURES

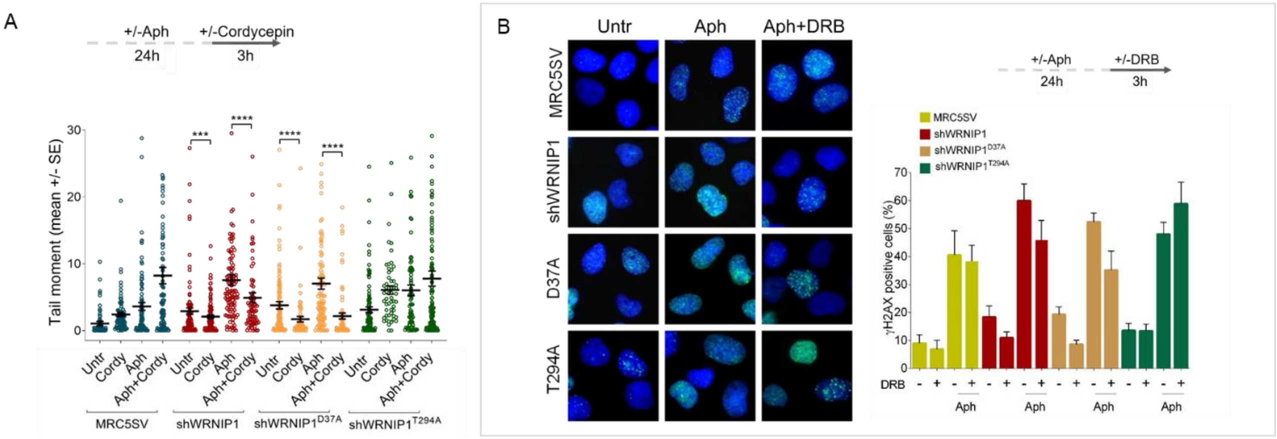


**Suppl. Figure 1**. **Loss of WRNIP1 or its UBZ domain increases DNA damage upon MRS**

(**A**) Evaluation of DNA damage accumulation by alkaline Comet assay. MRC5SV, shWRNIP1, shWRNIP1$^{D37A}$ and shWRNIP1$^{T294A}$ cells were treated according to the experimental scheme (0.4 µM Aph and 50 µM Cordy), and then subjected to Comet assay. Dot plot displays data as means from three independent experiments. Horizontal black lines represent the mean (***, $P < 0.001$; ****, $P < 0.0001$; two-tailed Student's t test).

(**B**) Analysis of DNA damage accumulation. MRC5SV, shWRNIP1, shWRNIP1$^{D37A}$ and shWRNIP1$^{T294A}$ cells were treated according to the experimental scheme (0.4 µM Aph and 50 µM DRB), and then immunostained for γ-H2AX. The graph displays data presented as percentage of γ-H2AX-positive cells ± SE from three independent experiments. Representative images of nuclei showing the different number of foci per nucleus are provided.

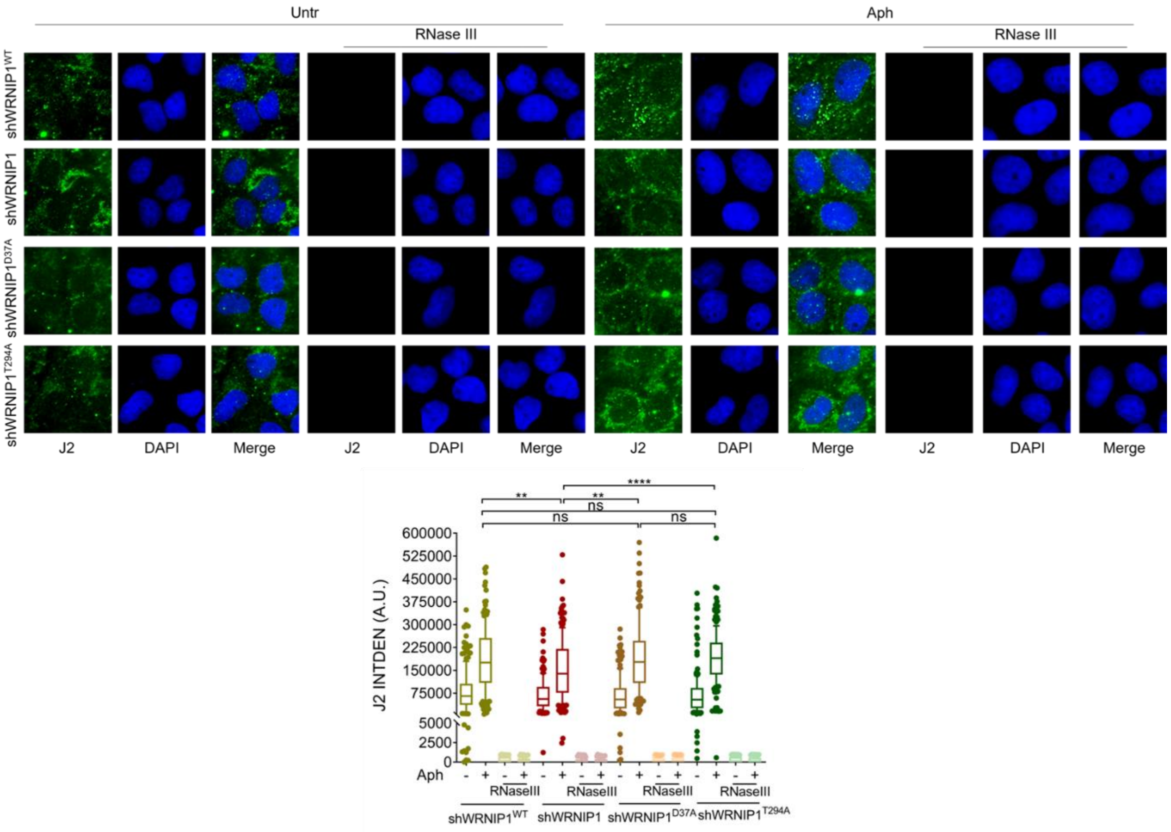


**Suppl. Figure 2. RNase III digestion significantly reduces the amount of dsRNA** Immunofluorescence analysis to determine dsRNA signal in MRC5SV, shWRNIP1, shWRNIP1$^{D37A}$ and shWRNIP1$^{T294A}$ cells treated or not with 0.4 μM Aph for 24 h. Cells were then fixed, subjected or not to RNase III digestion, and stained with anti-dsRNA monoclonal antibody J2. Representative images are provided for each single color channel. Nuclei were counterstained with DAPI. Dot plot shows nuclear dsRNA fluorescence intensity. The line represents the median value. Data are presented as means from three independent experiments. Horizontal black lines represent the mean. Error bars represent SE (ns, not significant; **, $P < 0.01$; ****, $P < 0.0001$; two-tailed Student's t test).

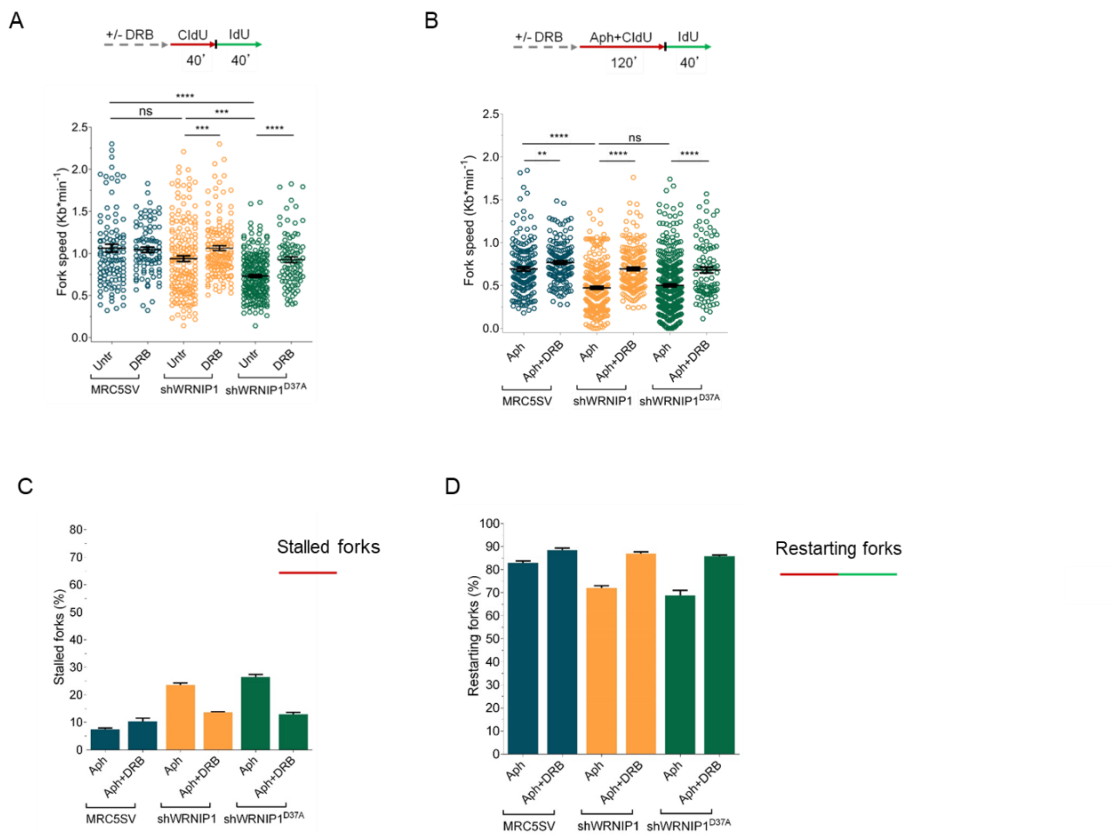


**Suppl. Figure 3**. **Transcription affects DNA replication in cells lacking WRNIP1 or its UBZ domain upon MRS**

Experimental scheme of dual labelling of DNA fibers in MRC5SV, shWRNIP1 and shWRNIP1$^{D37A}$ cells under unperturbed conditions (**A**) or upon MRS (**B**). After transcription inhibition by DRB, cells were pulse-labelled with CldU, treated or not with 0.4 µM Aph, and then subjected to a pulse-labelling with IdU. The graph displays the analysis of replication fork velocity (fork speed) in the cells. The length of the green tracks was measured. Mean values are represented as horizontal black lines (ns, not significant; **, *P* < 0.01; ***, *P* < 0.001; ****, *P* < 0.0001; Mann-Whitney test).

(**C** and **D**) The graph shows the percentage of red (CldU) tracts (stalled forks) or green (IdU) tracts (restarting forks) in the cells.

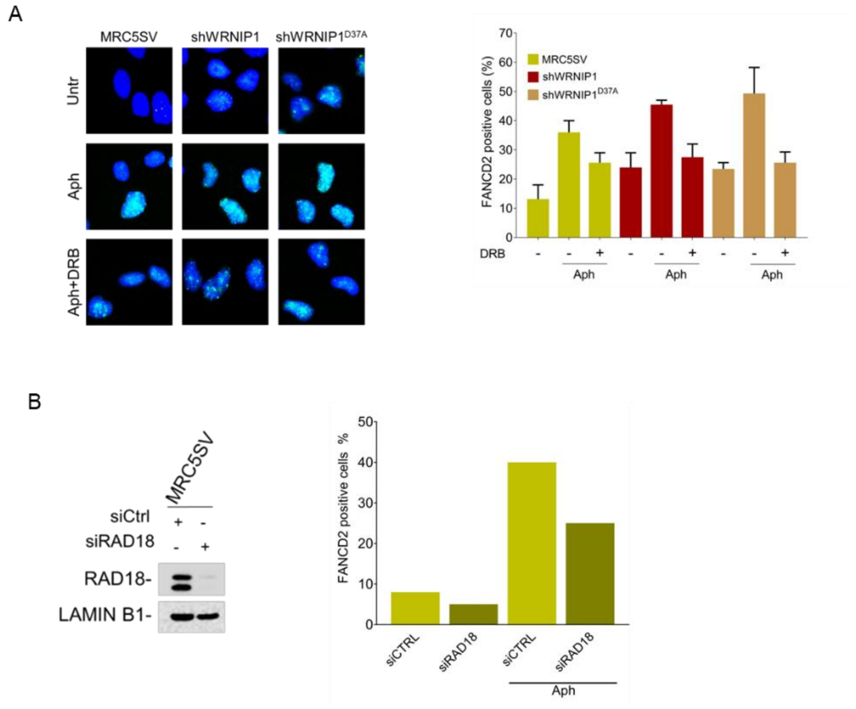


**Suppl. Figure 4**. **Loss of WRNIP1 or its UBZ domain results in FANCD2 pathway activation upon MRS**

(**A**) Evaluation of FANCD2 activation by immunofluorescence analysis in MRC5SV, shWRNIP1 and shWRNIP1$^{D37A}$ cells treated or not with 0.4 µM Aph for 24 h and 50 µM DRB for 3 h. The graph displays data presented as the percentage of FANCD2-positive cells ± SE from three independent experiments. Representative images of nuclei showing the different number of foci per nucleus are provided.

(**B**) Analysis of the dependency of FANCD2 activation on RAD18. MRC5SV cells were depleted of RAD18, and 48 h later, cells were fixed and immunostained for FANCD2. The graph shows data presented as the percentage of FANCD2-positive cells. Representative images of nuclei display the different number of foci per nucleus are provided. Western blot confirms RAD18 depletion in the cells. The membrane was probed with an anti-RAD18 and LAMIN B1 was used as a loading control.
